# Integrative Multi-Omics Approaches for Identifying Cervical Cancer Therapeutic Targets

**DOI:** 10.1101/2022.10.07.511244

**Authors:** Santosh Kumari Duppala, Rajesh Yadala, Aayushi Velingkar, Prashanth Suravajhala, Smita C Pawar, Sugunakar Vuree

## Abstract

After breast cancer, cervical cancer (CC) is one of the most common malignancies in women globally. Over 90% of chronic infections are caused by human papillomavirus (HPV) and its subtypes. Extensive research efforts are required to identify the treatment targets and prognostic indicators for recurring and metastatic cancers. It may be possible because of omics methods, including genomes, epigenomics, transcriptomics, proteomics, and metabolomics. High throughput (HT) data on the differential mRNA and miRNA expression and their crucial interrelationships enable promising integration and interpretation of the results. Clinical data and multi-omics have risen to the top of the heap in delivering molecular and cellular activities. They aid in comparing data from different omics approaches and bridging the gap between genotype and phenotype. Therefore, multi-omic techniques may improve the knowledge of the molecular basis of the physiology and primary cause of disease, revealing a new route for the prognosis, diagnosis, prevention, and therapy of human diseases.

## Introduction

Cervical Cancer (CC) is fourth cancer among all other cancers and is the second most cancer-causing in women. According to GLOBOCAN 2020, among 185 countries, the worldwide incidence rate is estimated at 604,127(3.1%), and the mortality rate is 341,831(3.4%), as shown by statistics (Sung et al. 2021). Despite breakthroughs in CC diagnosis, prognosis, and therapies, it remains the leading cause of death worldwide, particularly in developing nations. As a response, new biomarkers should be developed to identify novel biological markers in CC, boosting the chances of an early diagnosis with a favourable and even developing innovative and effective therapeutics based on these biomarkers.

HPV is a fundamental factor in CC development (Walboomers et al. 1999). There are numerous variants of HPV. HPV16 and HPV18 together are responsible for 71% of CC. It is the leading cause of cancer death in women, especially in low-income and developing countries such as Africa, Uganda, Sub-Saharan Africa, India, etc., compared to developed countries like Australia, New Zealand, and Northern America. CC is mainly caused by persistent infections of Oncogenic high-risk types of HPV and is one of the reasons but not a complete cause as other risk factors are also responsible such as reduced parity, multiple sexual partners, smoking, HIV, or chlamydia infections. CC is preventable after primary screening and Gardasil vaccination in girls below the age of 9-26yrs of age. CC shows strong evidence that HPV inactive cancers differ from HPV active cancers with the expression of oncogene E6/ E7, which drives to cause somatic mutations in CC.

Moreover, the drugs for CC with HPV active cancers and HPV inactive cancers show variation. For example, Gefitinib is more productive against HPV active CC as it is an EGFR receptor inhibitor. In the same way, Dasatinib is more effective against cervical HPV functional tumours (Banister et al., 2017).

While most HPV infections are transient and go by themselves within one or two years, persistent viral oncogenes E6 and E7 inactivate P53 and pRb, leading to genomic instability and integration of HPV to the host genome causing CC. Despite several advances in screening, treatment, and prevention of CC, the treatment modalities and effectiveness have not improved (Ouyang et al., 2020). Since the recurrence of CC with overall survival rate is low in most developing and underdeveloped countries. Therefore, identifying biomarkers and novel therapeutic targets for the survival and prognosis of the disease is vital. Moreover, in recent studies through TCGA, 70% of CC showed genomic alterations in PI3K/MAPK/TGFβ signalling pathways that offer clinical significance through therapeutic target agents. Various Multi-omics approaches were developed to untie the complexity of integrative biological systems.

Omics is a term used to comprehensively assess a set of molecules in biological science such as Genomics, proteomics, functional genomics, transcriptomics, metabolomics, etc. Multiomics approaches are HT technologies widely used in medical research that have enabled biological processes that involve highly effective interactive molecular approaches in the fields of genetics, epigenetics, Next-generation sequencing, etc., as DNA, RNA isolation, mRNA transcripts, protein, and metabolites play an important role in a corresponding biological system. These HT omics technologies have spearheaded the progress of systems biology era as a whole (Suravajhala et al., 2016). Therefore, multi-omic approaches increase the quality of understanding molecular compelling of the physiology and original cause of disease that shows the novel pathway for prognosis, diagnosis, prevention, and treatment of human diseases (Sun & Hu, 2016). Furthermore, the downstream analysis data generates statistical inference of data and associations depending on the parameters such as heterogeneity, sample size, and effect.

### Clinicopathology of CC

The most common CC are squamous cell carcinomas and adenocarcinomas, consisting of 70-90% and 12-25% of CC (Seoud, Tjalma, and Ronsse 2011; Vizcaino et al. 2000). In addition, a small proportion of CC (~35%) (Wright, Kurman, and Ferenczy, n.d.) represent adenosquamous carcinomas and other rare histological types, including neuroendocrine carcinomas (Gien, Beauchemin, and Thomas 2010). The cervix consists of two layers, the ectocervix, and the endocervix. The ectocervix is generally lined with non-keratinizing (SSE) stratified squamous epithelium and the endocervix with mucus-producing columnar epithelium. The cells in the squamocolumnar junction (SCJ), termed the transformation zone, are less stable and susceptible to viral infections (Waggoner 2003). In contrast, adenocarcinoma develops from the gland cells within the endocervical canal.

## Cervical precursor lesions Histologic and Cytologic classification

Cancerous lesions develop from precancerous lesions. Neoplastic adversity is determined by the intensity of the alterations, particularly on the epithelial layer. CIN is divided into three grades: CIN1, CIN2, and CIN3, which indicate that one-third, two-thirds, and almost the whole epithelial layer has abnormal cells, respectively (Solomon et al. 2002). The Pap and Bethesda systems are common cytological classifications. The European Pap classification is divided into five groups, PAP1 to PAP5, based on the severity of abnormal smears. (Glenn McCluggage 2013). The American Bethesda classification was created to distinguish between lesions that are likely to lead to CC “high-grade squamous intraepithelial lesions (HSIL),” and lesions improbable to progress “low-grade squamous intraepithelial lesions (LSIL)” (Park and Soslow 2009) the correspondence between the histologic and cytologic classifications are illustrated.

## Management of CC

Microinvasive CC can be treated by large loop excision of the transformation zone “LLETZ” or cone biopsy, depending on the (FIGO) stage. Radical hysterectomy with pelvic lymph node dissection or (chemo) radiation can be used to treat early-stage malignancies (Allam, Mohamed, et al. “2004). The most common treatment for advanced-stage malignancies is (chemo) radiation. Patients with stage IA cancers have a five-year viability rate of 99 to 100%, with 70-85% for stage IB1 and small IIa lesions, 50-70% for stages Ib2 and IIb, 29-50% for stage III, and 5-15% for stage IV13. As a result, early diagnosis is one of the most effective ways to save the lives of (CC) patients.

### Overview of Multi-Omics approach for diseases

The complexity of diseases depends on the nature of the data and samples. Genomes and proteomes play a vital role in complex biological networks, gene expression, and pathways. The isolation and product names of genes include RNA transcripts, cellular structure, transcription, and translational modification. The challenging step is understanding molecular mechanisms and information by studying multiple layers of omic data. Incorporating system approaches and multidimensional omics networks will address the research gaps and current knowledge of the tools. (Figure 1)

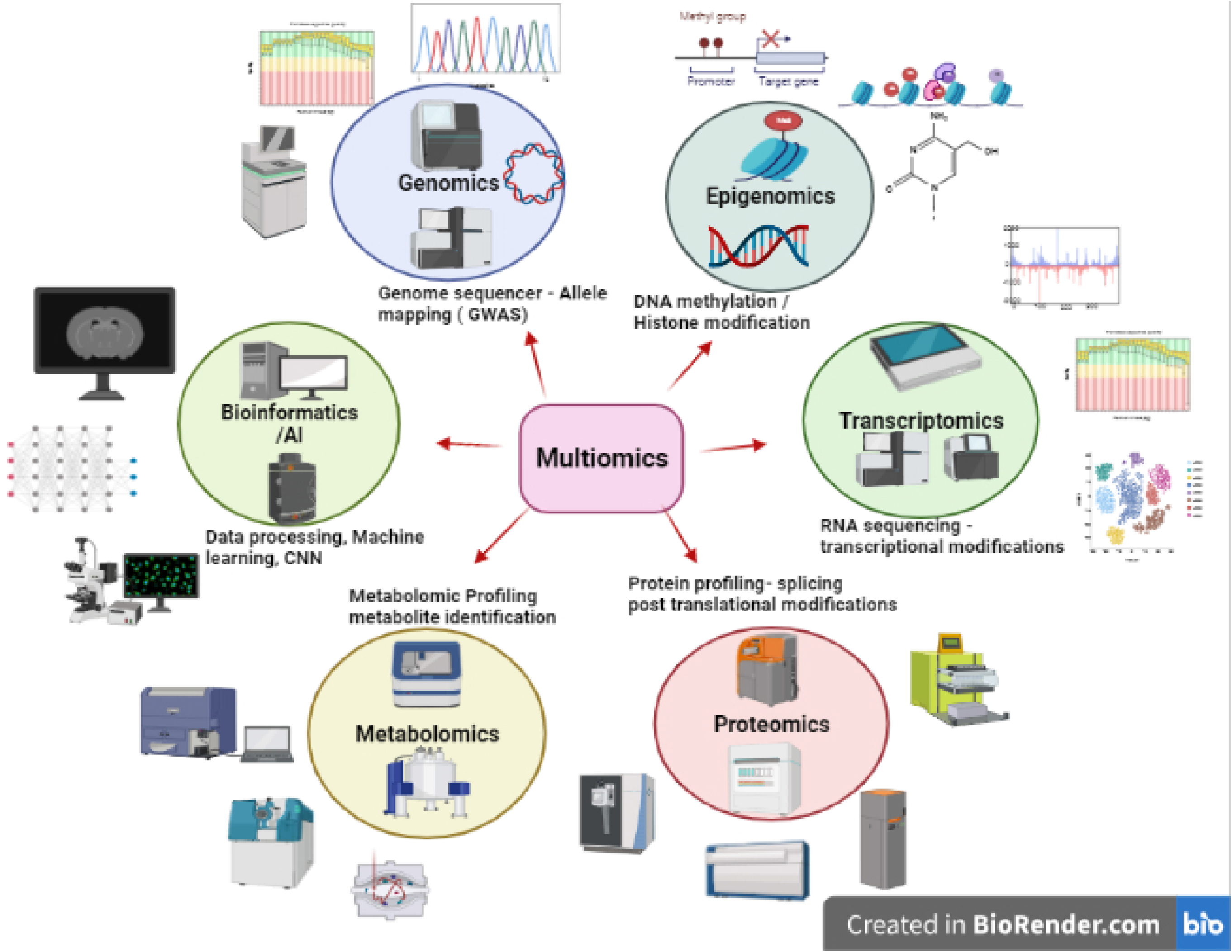

**Genomics studies in CC mainly focus** on recognizing genetic variants associated with the disease. Genome-Wide Association Studies (GWAS) play a vital role in identifying genetic variants, and it is related to complex phenotypic conditions (Hasin, Seldin, and Lusis, 2017). Moreover, the technologies include Next-generation sequencing, whole-genome sequencing, exome sequencing, genotype arrays, etc. Genome-wide association studies have identified the genes and genetic loci susceptibility in the development of human disease (Sun and Hu 2016) (Table 1). Genetic alterations play a role in the progression of CC. Integrated clustering of genomics and proteomics methods identified High Keratin squamous epithelium, low squamous epithelium, and Adenocarcinoma-rich clusters grouped by various HPV molecular characteristics. (Xu, Shen, and Xu, 2021) focused on studies relevant to the tumour microenvironment of CC and the differences between transcriptomic and somatic mutation studies through integrative multi-omics approaches based on TCGA.

**Table 1:** A table on Genomic Approaches in Cervical Cancer.

| **Type of Sequencing** | **Type & number of samples** | **Coverage & depth of sequencing platform** | **CNVs, SNVs,** | **Significantly Mutated Genes/Biomarkers** | **References** |
| --- | --- | --- | --- | --- | --- |
| Whole Exome | Frozen tissue tumor & blood - 178 Samples | The coverage is 47X for 178 samples |  | SHKBP1, ERBB3, CASP8, HLA-A TGFBR2, PIK3CA, EP300, FBXW7, HLA-B, PTEN, NFE2L2, ARID1A, KRAS and MAPK1 (HLA-A, HLA-B, NFE2L2, MAPK1, CASP8, SHKBP1, and TGFBR2 were found exclusively in squamous tumors) | (Burk et al., 2017) |
| Whole exome | Fresh frozen tissue - 54, Primary cell lines- 15 | Greater Depth and coverage of 173 independent reads per targeted base. | 56- CNV ERBB2/ | Somatic missense mutations – PIK3CA, ERBB2, GNAS, STK11/LKB1, t R135T and R135K, s R4007H, E4177K | (Zammataro et al., 2019) |
| Whole Exome and RNA Sequencing | 3 patient samples | 57X to 80 X |  | SYT2, BASP1, WI2-2373I1.2, JPH4, KRTAP9-238 1, HNRNPCL2, OR2L13, TAS2R46 64-somatic mutated genes | (Wu et al., 2019) |
| Sanger Sequencing | 103 tissue samples |  |  |  | (Tamura et al., 2018) |
| Whole-exome sequencing |  | (Illumina Hiseq) 2- X100bp |  |  | (Tamura et al., 2018) |
| RNA sequencing | Ect1, E6/E7 Cell lines<br>SiHa, MA 180 | Illumina HiSeq - 2500 |  | Gene fusion with FGFR3/TACC3 | (Tamura et al., 2018) |
| Whole exome sequencing | 70samples (including 25 CINs+10 blood samples) | 79.9× to 123× Illumina HiSeq X-Ten platform |  | *PIK3CA*, *ARHGAP5* and *ADGRB1* | Bao Chao Hai Et al. 2021 |

Searching for therapeutic targets from a Chinese woman’s study interrogates whole-exome sequencing data of 23 CC samples for neo epitopes and somatic mutation. In addition, a recent survey from cancer genome atlas long sequencing reads of HPV integration sites developed using ICC target genes.

#### Epigenomics

Epigenomics plays a central role in carcinogenesis; importantly, characteristic-wide DNA modifications like histone acetylation, DNA methylation, gene expression, and epigenome-wide association studies where the histone modifications are key regulators for gene transcription. These modifications play a vital role in the diagnostic and therapeutic targets. So, these epigenetic changes can be influenced by environmental factors, genetic factors, and long-lasting heritable changes, differentially methylated regions of DNA mostly used to detect metabolic syndrome, cardiovascular disease, and other cancer-causing genes that cause cancer; epigenetic modifications are tissue-specific and independent associated disease (Hasin, Seldin, and Lusis 2017). In a recent study, Histone deacetylases, DNA methyltransferases, and global genomic DNA methylation quantitation were performed by ELISA in quercetin to see the modulating alterations and promoter methylation in Hela cells of cervical tumour suppressor genes. Even molecular docking studies in quercetin work as a competitive inhibitor in DNA methyltransferases and histone deacetylates (Kedhari Sundaram et al., 2019). In an integrative survey, some important biomarkers of DNA methylation for cancer detection

## Transcriptomics

The study of the transcriptome in a particular biological cell, tissue, or organism at a particular moment or condition is known as transcriptomics. The term “transcriptome” describes the collection of all RNA transcripts from a gene in an organism. Transcription is a biological process that results in the production of RNA transcripts. Transcriptomics comprises everything related to RNAs, either coding or non-coding, including their function, location, transcription, expression, and degradation. Thus, transcriptomic studies are widely used to study differential gene expression and regulation pathways and hence have a vast significance in molecular biology and biomedical research, especially in disease diagnosis and profiling.

Transcriptomic analysis benefits in detecting genes dysregulated in cancer cells by comparing the transcriptomes of the tumour and non-tumour tissues, which gives the essential information regarding dysregulated genes in tumours, making it a beneficial technique to identify biological markers of cancer progression and disease severity. Apart from gene expression and regulation studies, transcriptomics also plays a vital role in identifying gene changes and mutations.

One deregulated gene identified is a bone gamma-carboxyglutamic acid-containing protein (BGLAP). Also recognized as osteocalcin, it is an 11-kDa protein hormone produced and released by osteoblasts. The major functions of BGLAP are regulating bone matrix mineralization and controlling blood coagulation. However, the transcriptomic studies made by (Bettadapura, Munivenkatappa, and Madhunapantula 2018) indicated that cervical carcinomas show higher levels of BGLAP expression. Thus, their studies concluded that BGLAP upregulation is seen only in cancer tissues and not in normal tissues, indicating that it might be a useful therapeutic target in this malignancy.

The study made by (Kori and Arga 2018) led to the identification of cervical transcriptomic codes of CC using 5 distinct transcriptome datasets. They have found 113 down-regulated genes and 199 upregulated genes in CC. These dysregulated core genes encode proteins involved in hormones, signalling molecules, modulators, enzymes, receptors, transporters, structural proteins, etc. The down regulated proteins include KAT2B (lysine acetyltransferase enzyme), ESR1 (estrogen receptor), WNK1 (serine/threonine-protein kinase), and FGFR2 (fibroblast growth factor receptor 2). In contrast, upregulated proteins are GSK3B (glycogen synthase kinase 3 beta), PARP1 (poly ADP-Ribose polymerase), PCNA (proliferating cell nuclear antigen), and CDK1 (cyclin-dependent kinase 1). In addition, their studies found that several deregulated genes are involved in metabolism pathways in CC by combining useful transcriptomic data with the genome-scale metabolic network. Their report concluded that the arachidonic acid metabolism pathway was one of the prospective therapeutic targets, and the biomarkers indicated above could be employed for CC screening or treatment.

Another reason for CC is variations in the gene expression responsible for causing cancer. Micro RNAs or miRNAs play a positive role in gene expression regulation. MicroRNAs are a small class of single-stranded non-coding RNAs that mainly bind to the 3’ untranslated regions (3’ UTRs) of the mRNA and negatively control gene expression or sometimes completely repress the mRNA. In most cancer tissues or cells, either oncogenic miRNAs are upregulated, or tumour suppressor miRNAs are downregulated, thereby endorsing the expression of cancer. In these conditions, transcriptome analysis utilizing RNA-sequencing (RNA-seq) has emerged as a powerful method for studying gene expression patterns. In CC cells, about 250 miRNAs are differentially expressed. One of them is miRNA-214 or miR-214, downregulated in all three stages of cervical intraepithelial neoplasia (CIN) (Yeung et al. 2017).

(Sen et al., 2020) used transcriptomic analysis to assess gene expression levels of several genes in CC cells following miR-214 up and downregulation. They used C33A and CaSki cell lines. The Knockdown of miR-214 was done using CRISPR/Cas 9 technique to downregulate the expression. The over-expression of miR-214 was done by pCDNA-HisA plasmid transfection, and gene expression was checked by RNA extraction followed by quantitative RT-PCR in both cases. Their studies have concluded that increased levels of apoptosis are observed during overexpression of miR-214 and reverse in case of knockout. As a result, these genes too could be employed as possible biomarkers to describe tumour prognosis in various cancer stages, including CC.

Using miRNA microarray, (J. H. Li et al. 2011) described several variably expressed miRNAs in CSCC and adjacent non-cancerous tissues. The study included 1145 miRNAs and found that 7 of them, including miR-886-5p, varied substantially between tumour and non-tumour tissues. miRNA microarray and bioinformatics analysis revealed that HPGD (15-hydroxyprostaglandin dehydrogenase), which can limit cell proliferation and migration, is a direct target of miR-146b-3p in CC (Yao et al. 2018).

Another type of RNA that plays a vibrant role in regulating transcription factors is Long non-coding RNAs (lncRNAs). These are non-protein-coding, functional RNA molecules with more than 200 nucleotides. The combination of lncRNA and mRNA microarray studies by Liu et al. in 2018 identified 1621 lncRNAs and 1345 mRNAs expressed differentially between high and low-risk squamous CC. Another group of researchers (J. Huang et al., 2018) used an HT microarray to compare lncRNAs and mRNAs expression profiles in cancerous and normal tissue samples in the same year. They found that 5844 lncRNAs and 4436 mRNAs were differentially expressed in CC compared to normal tissues.

LncRNA microarray experiments proved that lncRNAs play a key role in regulating malignant genes by promoting or suppressing them. For example (Jing et al. 2015) conducted their studies on lncRNA named HOX Antisense Intergenic RNA (HOTAIR) because of its substantial role in regulating the expression of many oncogenic genes. They detected the HOTAIR expression in 4 CC cell lines (CaSki, C33A, HeLa, and SiHa cells) and found that HeLa and SiHa presented a much higher level of HOTAIR compared to CaSki and C33A cells. Furthermore, they discovered that the lncRNA HOTAIR inhibits p21 in CC, promoting cell cycle progression, proliferation, migration, and invasion hence serving as an oncogene. Therefore, its overexpression leads to CC. By contrast, lncRNA-LET acts as a tumour suppressor, and its down-regulation leads to CC. Research by (Jiang, Wang, and Yang 2015) revealed that patients with CC who have down regulated lncRNA-LET have a significantly lower overall survival rate than those who have upregulated lncRNA-LET.

Comprehensive lncRNA profiling analysis by (W.-Y. Zhang et al. 2019) was utilized to screen lncRNA expressed differently in CC. They reported two lncRNAs in two articles, which are highly expressed in CC. They are lncRNA NCK1-AS1 and lncRNA ANRIL, upregulated in CC tissues. Downregulation of ANRIL inhibited cell migration, proliferation, and invasion in CC cells. Sun et al., using lncRNA microarray, revealed that 4 circulating lncRNAs, PVT1, HOTAIR, AL592284.1, and XLOC 000303 included, might be promising biomarkers for CC prediction (Table 2).

**Table 2:**
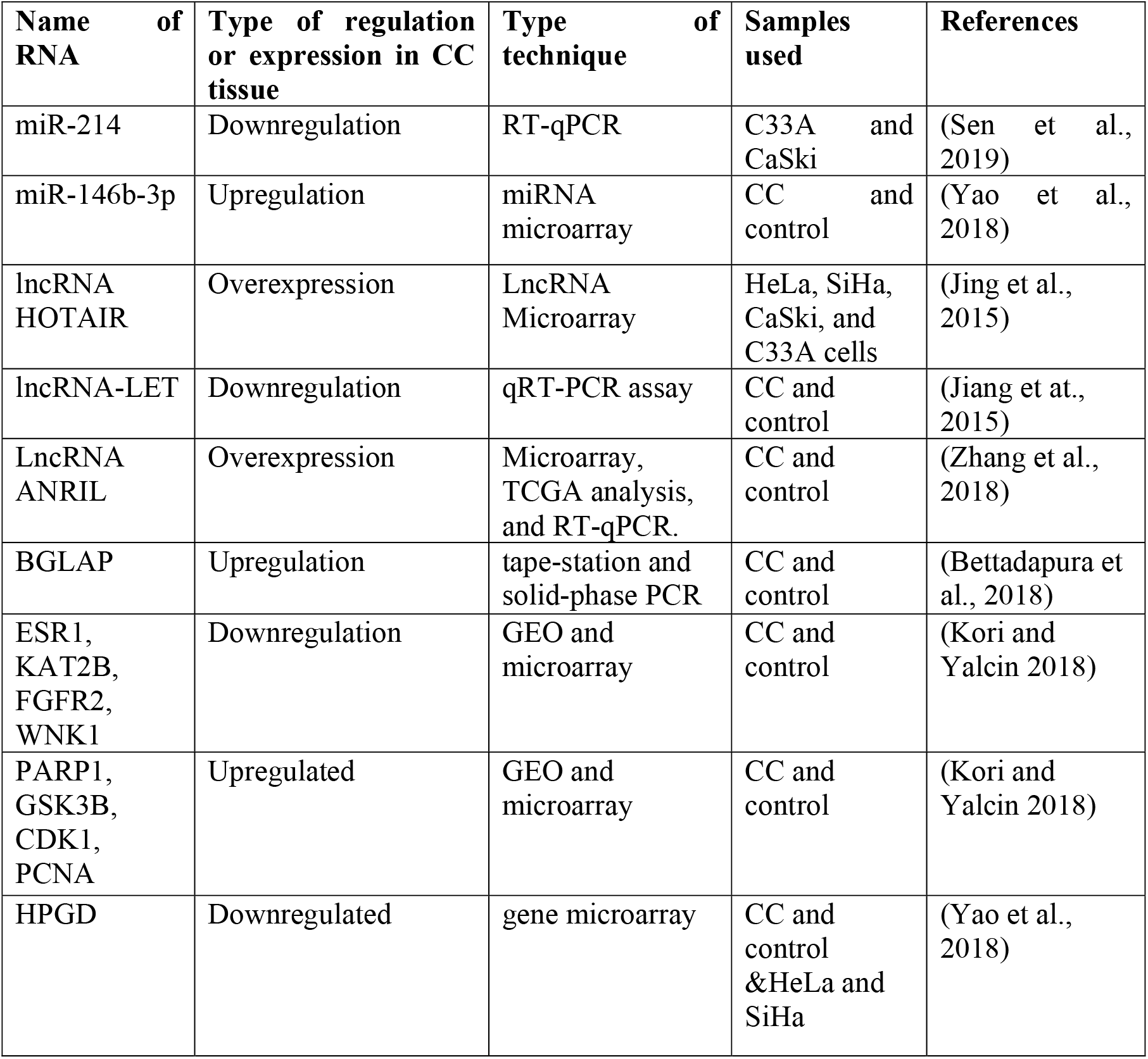
A table on Transcriptomics techniques used in Cervical Cancer.

Wallbillich et al., 2020, identified 42 genes as being crucial for squamous cell carcinoma of the cervix patients’ survival (SCCC). Most of these genes are associated with the key features of cancer cells, like cellular proliferation, migration or invasion, and/or metastasis. They used the TCGA dataset for this study. It has the largest number of patients, and the gene expression data is of the highest quality for any publicly available dataset of patients with CC. The gene expression studies using transcriptomics revealed that the increased expression level of 42 genes led to a gradual decrease in survival rate. Machine learning recognized a 9-gene signature that appears adequately accurate to predict survival. Transcriptomic risk score (TRS) for mortality was computed for these 9 essential genes. Among TCGA SCCC patients analyzed, the survival prediction worsened either with the increased expression level for each high-risk gene or a greater number of those genes with high expression levels in a patient’s tumour or by both. Thus, the transcriptomic studies of HPV help deeply understand the connection between the oncoproteins of HPV and CC.

Chen et al. (2019) worked on next-generation sequencing of HPV to understand the transcriptional level differential expressions in HPV-induced cancers. Their research focused on the best-known human CC SiHa cell line obtained from the American Type Culture Collection (ATCC). The cells are cultured at 37°C and 5% CO2 in DMEM (Corning, 10-013) or RPMI 1640 (Corning, 10-040-CVR) media mixed with 10% foetal bovine serum (FBS). TRIzol reagent was used to extract total RNA, and the PrimeScript RT reagent Kit synthesized cDNA. A HiSeq 2500 platform was used to sequence the transcriptome. They looked at 6 normal cervical tissues without HPV infection, 6 CC tissues with HPV16 disease, and SiHa cells with or without 16E6/E7 knockdown using HT RNA deep-sequencing techniques. They have identified 1347 and 586 differential expressed genes in CC tissues or cells. In addition, they have found 140DEGs in both sets of sequencing data. Their transcriptomic studies or transcriptional factor analysis in TransFind, TF2DNA database, and expression correlative analysis suggested that transcription factors, including E2F8 and EGR2, may play critical roles in CC associated with HPV.

Combining metabolomics and transcriptomic analysis has recently shown a significant progression in CC research. Metabolomics is widely used to understand cancer metabolism and identify biomarkers to infer the onset and progression of cancer. (Yang et al., 2017) used metabolomic profiling with transcriptomics data to validate the potential diagnostic biomarkers of CC. They selected five metabolites as CC biomarker candidates, which had a brilliant performance in distinguishing between CC and normal. It has been proven to be a viable approach for CC screening and diagnosis. This study confirmed combinational analysis of metabolomics and transcriptomics as promising methods to examine the mechanism of carcinogenesis and discover more consistent biomarkers.

## Proteomics in CC

The proteome is a term that refers to all proteins that can be investigated after they have been produced through alternative splicing and post-translational modifications (PTMs). As a result, proteomics research uncovers changes in PTMs and protein expression levels critical for protein function regulation. The primary purpose of proteomics is to better understand biological systems by examining the proteome that makes up a cell. For example, there have been genetic and proteomic changes in cancerous conditions, particularly in the pathways involved in cell development, proliferation, and apoptosis signalling. The most significant changes include (1) mutations in DNA replication and repair genes, (2) mutational transition of proto-oncogenes into oncogenes, and (3) mutations that suppress the activity of tumour suppressor genes. These mutations affect all somatic and germ cells, which may be genetic or irregular. Proteomics is a quick, sensitive method with extensive proteome coverage that uses expression, structural, and functional proteomics to decode changes in all of the above proteins.

The proteome of CC has been studied in several biological samples like fresh tissue, CC cell lines, formalin-fixed paraffin mediated (FFPE) tissue, cervical, vaginal fluid (CVF), serum, plasma, and urine. Proteomics research in CC is primarily conducted on tissue samples, either fresh or FFPE, depending on availability. The disadvantage of using FFPE samples is that the proteins may be damaged during the sample preparation procedures.

(Bae et al. 2005) compared healthy tissue proteins with CC biopsy protein patterns by MALDI-TOF and 2-DE. As a result, he discovered DE of 35 proteins, 17 of which were up-regulated and 18 down-regulated. Among all, twelve proteins were previously known to be involved in tumour development (annexin A2 and A5, pigment epithelium-derived factor, keratin 19 and 20, smooth muscle protein 22 alpha, α-enolase, heat shock protein 27, squamous cell carcinoma antigen 1 and 2, apolipoprotein a1, glutathione S-transferase). In comparison, 21 proteins were discovered for the first time in this study. At the same time, other proteins, including PI-TP-α, 60-kDa SS-A/Ro alternative proteins, Rho GDP dissociation inhibitor beta protein, and src homology 3 (SH3) domain-containing protein HIP-55, were newly identified.

The abundance of proteins found in proteomics studies may hide the presence of significant but low concentrated protein levels. For example, (Gu et al. 2007) published proteomic study results showing substantial levels of cervical dysplasia. More than two hundred proteins in tumour cells were identified using LC-MS and Laser Capture Microdissection. Furthermore, impaired to normal cervical epithelial cells, samples of high-grade squamous intraepithelial lesions showed a considerable up-regulation of nuclear and mitochondrial proteins.

(Zhu et al. 2009) used MALDI-TOF MS and 2D DIGE to investigate variations in the expression of zinc-finger protein 217 and Tyk2, S100A9 in squamous CC and concluded that these proteins could be beneficial in diagnosis and therapy.

(W. Wang et al. 2014) investigated three proteins involved in CC metastasis processes: FABP5, HspB1, and MnSOD, in 2014. They, too, used MALDI-TOF/TOF MS and 2D-DIGE to examine tissue samples with and without pelvic lymph node metastases. (Zhao et al., 2015) presented their findings by utilizing 2D-DIGE and MALDI-TOF/TOF MS to identify proteins that vary in expression between a healthy cervix and cervical intraepithelial neoplasia (CIN) and cervical squamous cell carcinoma (CSCC). Western Blot (WB) and immunohistochemistry were used to confirm the findings. The S100A9 protein was up-regulated significantly, while Eukaryotic elongation factor 1-alpha-1 (eEF1A1) was down-regulated in all three samples. S100A9 was localized in the cytoplasm, with a positive expression rate of 100% in CSCC, 70% in CIN, and 20% in the normal cervix, according to IHC data. On the other hand, eEF1A1 was mostly found in the cell plasma with positive expression in 60% of CSCC, 73.3% of CIN, and 70% of the normal cervix. PKM2 was localized in cell nuclei, with positive expression rates of 75% in CSCC, 93.3% in CIN, and 100% in the healthy cervix, with statistically significant differences. By the author’s findings, these 3 proteins might be prospective biomarkers for early CC diagnosis and novel treatment targets (Table 3).

**Table 3:**
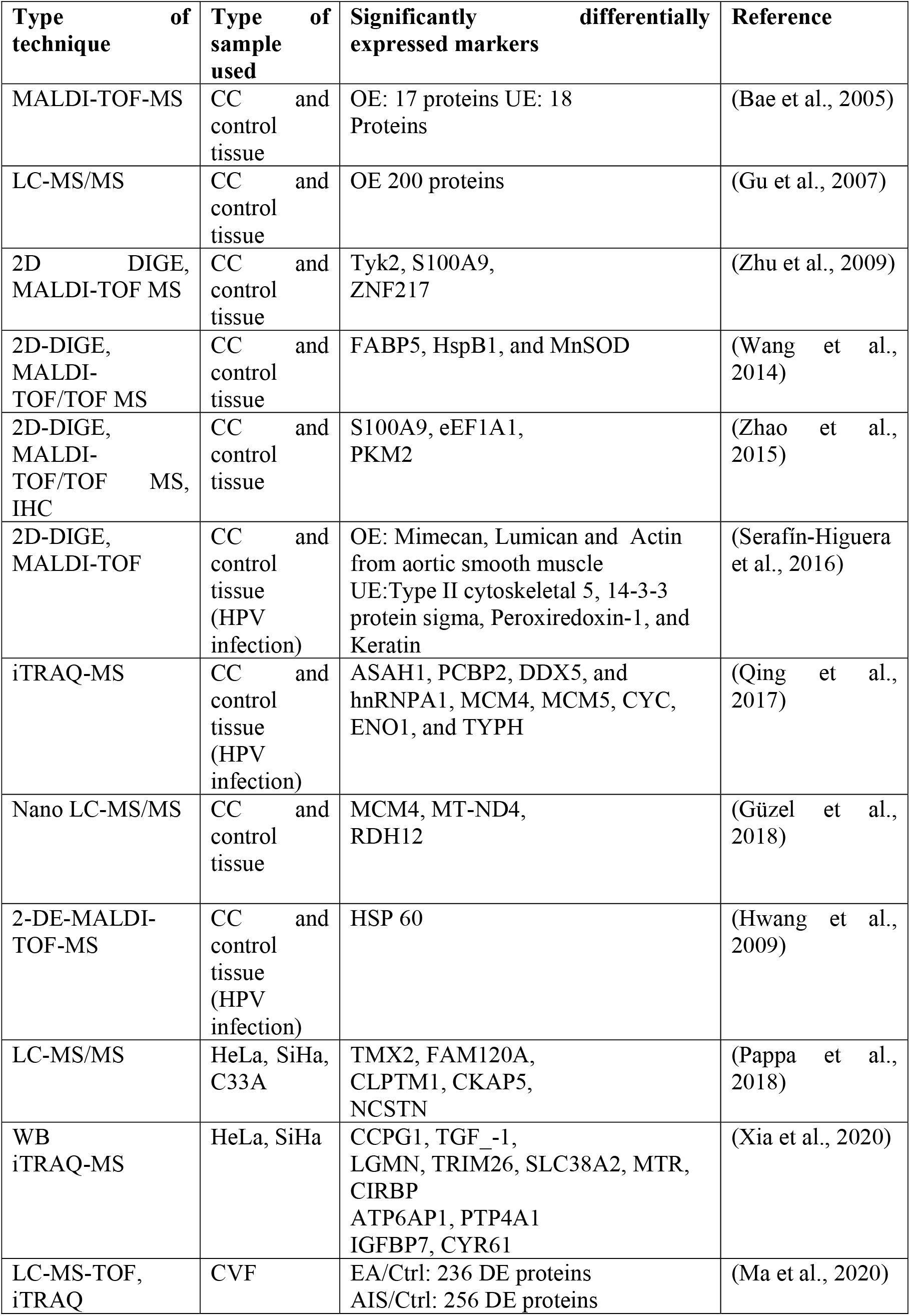
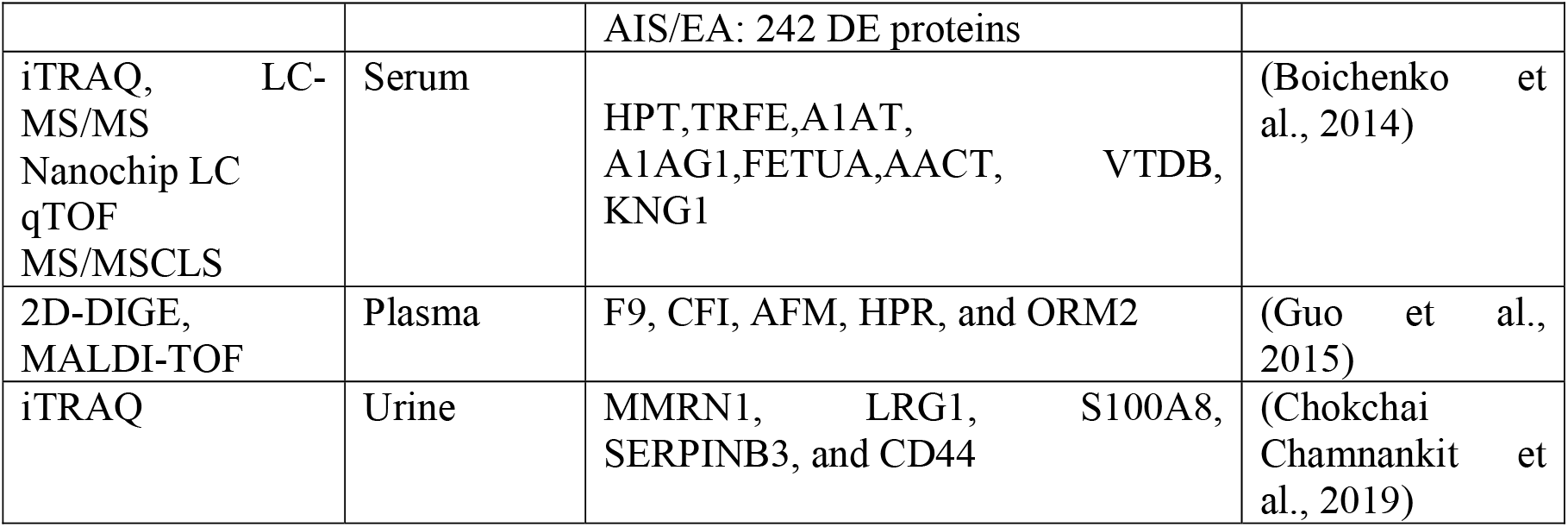
A table on Proteomics and techniques used in Cervical cancer.

(Hwang et al. 2009), by 2-DE-MALDI-TOF-MS, decoded 19 DEPs in CC caused by HPV infection, among all expressions of Heat Shock Protein 60 (HSP 60) is highly upregulated in CC. He also validated the upregulation of HSP60 by other techniques as an identifier of CC prognostication.

Furthermore, (Serafín-Higuera et al. 2016) used MALDI-TOF and 2D-DIGE to compare 6 cervical HPV-16 and 4 surgical specimens without lesions or traces of HPV infection associated with CC. The findings demonstrate that Mimecan, Lumican, and Actin from the aortic smooth muscle showed increased CC expression. In contrast, type II cytoskeletal 5, keratin, 14-3-3 protein sigma, and peroxiredoxin-1 decreased protein expression. Therefore, the authors concluded that these proteins could act as biomarkers in CC diagnosis and treatment.

(Qing et al., 2017) used iTRAQ-based proteome analysis to find DEPs in HPV-infected cervical squamous cell carcinoma patients and healthy controls. The study reveals that PCBP2, ASAH1, hnRNPA1, DDX5, MCM5 and 4, ENO1, TYPH, and CYC increase during cervical carcinogenesis, which may be linked to HPV infection.

Research on normal healthy tissue and CC tissue using laser capture microdissection and high-resolution mass spectrometry (Güzel et al., 2018). As a result, he identified a significant increase in MCM family proteins’ expression, which plays a key role in DNA replication. Thus, these proteins can be exploited as therapeutic targets in the early stages of CC.

The research was conducted in CC-related cell lines and proteome analysis in CC tissues. Membrane proteomics of CC cell lines reveals that many proteins were expressed differently when cancer cell lines (HeLa, C33A, and SiHa) were compared to the normal HCK1T cell line. A constitutive coactivator of PPAR-like protein 1, thioredoxin-related transmembrane protein 2, cleft lip and palate transmembrane protein 1, cytoskeleton-associated protein 5, and nicastrin are among the most notable membrane proteins that were upregulated in all three cancer cell lines, regardless of the presence of HPV infection (Pappa et al. 2018).

Metformin, an anti-diabetic drug, has been found to suppress tumour growth in recent years; however, the mechanism is unknown. iTRAQ-based quantified proteomic study of the inhibition of CC cell invasion by Metformin showed 53 differentially expressed proteins wherein most of them were related to apoptosis. It inhibits the proliferation and invasion of CC cells via regulating the insulin signalling system and interfering with cell proliferation and apoptosis (Xia et al., 2020).

Cervical-vaginal fluid (CVF) is an inconsistent biological fluid used to research and find biomarkers for the early detection of CC. The major advantage of identifying biomarkers from CVF is that most proteins are localized either in the extracellular region or the cytoplasm. A recent study (Ma et al. 2020) includes proteome analysis of endocervical and cervical adenocarcinoma in situ (EA and AIS, respectively) with controls. EA/Ctrl comparison found 237 differentially expressed proteins, AIS/Ctrl comparison found 256 differently expressed proteins, and EA/AIS comparison found 242 differently expressed proteins. GO studies of all the proteins reveal heme protein myeloperoxidase and APOA1 as significant biomarkers in CVF fluids.

In most cases, cancer markers circulate within the bloodstream at a concentration of ng/mL. By comparing the profiles of plasma and serum samples of healthy controls with that of CC patients, powerful methods such as 2-DE gels paired with MS identification can be used to characterize these proteins. Early detection of CC can be aided by identifying serum indicators. (Boichenko et al., 2014) investigated the proteome of serum obtained from CIN and early CC against controls by iTRAQ labelling followed by LC QTOF MS/MSCLS and LC-MS/MS. The HPT, A1AT, TRFE, FETUA, A1AG1, AACT, KNG1, and VTDB were identified as possible markers in the first investigation. They tested the selectivity of the identified biological markers for CC by examining sera from late-stage ovarian cancer patients. After their research, the authors found that HPT, A1AG1, and A1AT are not specific to CC. (Guo et al., 2015) used MALDI-TOF MS and 2D-DIGE gels to assess early-stage CC samples. Following the analysis, ten proteins were identified as potential CC biomarkers. The proteins studied are involved in lipid metabolism and molecular transport networks. Five novel protein plasma biomarkers (F9, CFI, AFM, HPR, and ORM2) for cervical precancerous lesions and prognostic evaluation of CC were discovered during the screening of prospective plasma protein biomarkers for the transition of cervical precancerous lesions into cervical carcinoma.

In most circumstances, urine analysis is a good non-invasive tool for easily detecting biomarkers. For example, urinary biomarkers for CC diagnosis by quantitative label-free MS analysis reveal MMRN1, LRG1, S100A8, SERPINB3, and CD44 as potential markers for early prognosis of CC (Chokchaichamnankit et al. 2019).

## Metabolomics

Metabolomics is quickly gaining traction in medicine due to its enormous potential for developing diagnostic and prognostic tools for tracking illness progression and determining the effects of therapeutic drugs.

(Wishart 2016). Furthermore, metabolomic profiling can aid in deciphering diseased molecular pathways, thus allowing for the discovery of novel therapeutic targets (Wishart 2019). Metabolomics has been broadly applied to infer the genesis and development of cancer through cancer metabolism and biomarker identification (B. Wang et al., 2015; P. Yin and Xu, 2013). Metabolites are the by-products of various biological processes, and they can serve as particular biomarkers for biological events like genetic variations and epigenetic perturbations (Nicholson and Lindon 2008). Under this, the unmet demand for improved diagnosis and personalized therapy can be addressed through metabolomics.

Identifying biomarkers for CC has ushered in a new age distinguished by advances in molecular biology, proteomics, and metabolomics (P. Shi, Zhang, and Ye, 2015; Walker et al., 2017). The final products of numerous biological activities are metabolites. They show promise as reliable biomarkers by detecting various upstream biological processes (Khatami et al., 2019; Nicholson and Lindon, 2008). It has been utilized to thoroughly investigate and quantify metabolites in biological systems using techniques including mass spectrometry (MS) or spectroscopy and nuclear magnetic resonance (NMR) paired with chromatography (Tebani et al. 2016; Wishart 2008). Metabolomics has proven to be an effective technique for detecting metabolic alterations in tumour formation and development and identifying and developing non-intrusive biomarkers for cancer prediction, diagnosis, and monitoring in cancer research (López-López et al., 2018; F. Zhang et al., 2017). Compared to normal (healthy) controls, changes in metabolic pathways and their profiles can help better understand dysregulated metabolism in tumour prediction and development (S. Huang et al. 2016).

Hou et al.’s study presented the metabolomics paradigm for predicting the chemotherapeutic response for CC sufferers before non-adjuvant chemotherapy (NACT) (Hou et al. 2014). This study used ultra-performance liquid chromatography coupled to time-of-flight mass spectrometry (TOFMS), which has proven to be a powerful technique with high specificity and sensitivity. By its application, one steroid, one sphingolipid, and two amino acids (aa) were highly correlated with the complete pathological response (pCR). Despite the variability of histopathological and clinical features among individuals in each response class, plasma samples may still be used to separate them into different clusters.

Multivariate analysis was used to determine if these biomarkers can predict post-therapy responses separately by evaluating the effect of pathological parameters for such biomarkers. These biomarkers were found to be independent predictors of chemotherapy treatment response later.

The sensitivity and specificity of the prediction model created using the chosen model are high. Given the difficulties in predicting chemotherapy response in cancer patients worldwide, this future metabolomics research may make a new window for individuals to pick the most promising medication or even genuinely “tailored treatment” in clinical practice. Notably, the chosen biomarkers are substantially linked with the chemotherapeutic response, as shown by the multivariate regression analysis between post-therapy responses and biomarkers and clinical characteristics indicators. Although altered metabolic pathways connected to identified biomarkers are highly linked to treatment responses, bigger cohort studies are needed to corroborate these findings and establish metabolic linkages with pathologically different subtypes, not simply squamous CC.

Further, the 2 biomarkers discovered using LC-MS analysis performed exceptionally well in prediction. The metabolites provided more cellular metabolism information and solid prototypes that could accurately predict chemotherapeutic response strategies when authenticated in a relatively large subset of patients. The high accuracy of the metabolomics technique in classifying plasma metabolites implies that it could be particularly useful for predicting NACT response in CC patients worldwide.

Since cancer patients’ metabolic processes differ from healthy people (T. Zhang et al. 2013; 2012; H. Zhang et al. 2012; Xie et al. 2012), any alterations in metabolism might indicate metabolic instability.

Hou and his team used R package CAMERA to annotate isotope peaks after converting the raw data to identify and discover metabolic biomarkers in CC patients. As a result, 562 heights (variables) were determined based on the identification verification using relevant raw original data and the labelling of peaks produced using CAMERA. Their research team used the Wilcoxon Kruskal-Wallis test and multivariate analysis to find a set of metabolites with the highest individual potential to predict responsiveness before chemotherapy. The p0.05 and VIP1 criteria were used to select the candidate biomarkers. 2 out of the 4 biomarkers (L-valine and L-tryptophan) were verified with an external independent standard reference.

They discovered that the metabolites mentioned above were engaged in the pathways of tryptophan, amino acid, and lipid metabolism after mapping all distinct metabolites into the KEGG database. These findings revealed a link between CC and metabolic abnormalities.

The primary aim of the studies in CC was to find new possibilities for early tumour diagnosis or to develop prognostic frameworks. Therefore, metabolomic profiles of cancer patients (blood, urine, tissue samples, and cervicovaginal lavage) were analyzed and differentiated from those of healthy women and those with benign uterine diseases (e.g., precancerous cervical intraepithelial neoplasia [CIN], cervicitis (Yang et al. 2017; Abudula et al. 2020; Khan et al. 2019; Ilhan et al. 2019; Tokareva et al. 2020; M. zhu Yin et al. 2016; Tokarz, Adamski, and Rižner 2020). As a result, algorithms with high diagnostic properties have been reported (Yang et al., 2017; Abudula et al., 2020; M. zhu Yin et al., 2016).

With a sensitivity of 93%, specificity of 91%, and AUC of 0.97 (M. zhu Yin et al. 2016) revealed the identification and verification of a model that included a combination of 2 phosphatidylcholines, 4 metabolites, and 2 lysophosphatidylcholine along with demonstrating discrimination between CC patients and individuals with uterine fibroids. With an AUC of 0.99, (Yang et al. 2017) presented a diagnostic technique that discriminated CC patients from healthy controls. 3 long-chain fatty-acid derivatives, lysophosphatidylcholine and bilirubin, were utilized in this approach. (Table 4)

**Table 4:** A table on Metabolomics AUC, SE, SP used in Cervical cancer by. Yang et al., 2017

| Biomarker | AUC | SE | SP |
| --- | --- | --- | --- |
| Bilirubin | 0.88 | 0.91 | 0.71 |
| LysoPC (17:0) | 0.94 | 0.94 | 0.86 |
| N-oleoyl threonine | 0.85 | 0.83 | 0.79 |
| 12-Hydroxydodecanoic acid | 0.92 | 0.94 | 0.79 |
| Tetracosahexanoic acid | 0.82 | 0.75 | 0.76 |
| Combination | 0.99 | 0.98 | 0.99 |

Further, (Khan et al. 2019) developed a diagnostic algorithm based on metabolic profiling in plasma samples that demonstrated strong diagnostic qualities for separating healthy women from patients with CIN (AUC of 0.82) and healthy women from patients with CC (AUC of 0.83)

(Zhou et al., 2019) on the other hand, a studied plasma metabolome from CC patients without and with recurrence presented predictive algorithms with AUC > 0.9. Metabolites also distinguish between low and high-grade squamous intraepithelial lesion patients and HPV-negative healthy premenopausal women with AUC > 0.8. Patients with invasive CC and healthy women have AUC of 0.92 (Ilhan et al., 2019). In two examinations of cervical tissue samples, 17 metabolites were discovered that might be utilized to distinguish between CIN and normal cervical tissue or squamous CC and between squamous CC and CIN (Abudula et al., 2020; Tokareva et al., 2020) revealed that nonpolar glycerolipids differentiate cervical boundary tissue from CC tissues with an AUC of 0.95 using shotgun metabolomics.

Several investigations found different quantities of aa in plasma or serum for CC, though the results were inconsistent. In plasma, Hasim et al. found reduced valine, isoleucine, alanine, histidine, glycine, and glutamine, but Ye et al. found greater levels of alanine and glycine in serum (Tokarz, Adamski, and Rižner 2020). Although these metabolites are the basic units for tumours, biomass levels in the blood are aberrant in cancer patients (Wishart 2019). Lactate levels in cervical carcinoma tissues are greater (Abudula et al. 2020), which may help tumour formation. The Warburg effect (Wishart 2019) describes how tumour cells use aerobic glycolysis to create ATP, with lactate as the final result. Lactate, produced by tumour cells, has an angiogenic effect (Murray and Wilson 2001; Wishart 2019) and promotes cancer growth.

Various other studies have revealed alterations in single combinations of metabolites in individuals with CC. Despite this, all the changes mentioned are too comprehensively spread to be interconnected on the metabolic pathway map.

Although the generated models were statistically confirmed, any absolute model for these new biomarkers must be empirically tested. Only one study gathered a second batch to evaluate the biological markers discovered in the first cohort (Khan et al., 2019), indicating that almost all of the investigations included were in the early stages of biomarker discovery.

A deeper analysis will determine if metabolic biomarkers can distinguish between endometrial and CC and other cancer types. It is critical because, among other metabolic alterations, all cancer types modify their metabolism to depend more on glucose and consume more (Hanahan and Weinberg 2011). As a result, any blood-based biomarkers distinctive of CC or endometrial cancer must be thoroughly verified to ensure that they are not false positives. One of the most difficult issues is establishing a reliable distinction between various diseases through biomarker identification.

Some metabolomics research has already been used in CC studies because altered metabolites and pathways might better understand dysregulated metabolism in tumorigenesis and progression (Garbett et al., 2014; Chai et al., 2015; Liang et al., 2014; Ye, Liu, and Shi 2015).

Hasim et al., for example, profiled 19 aa for CC (Bjerrum et al. 2014), while Yin and his team detected four lipids as novel CC biomarkers (M. zhu Yin et al. 2016). However, because the sample sizes in these studies were comparatively smaller, the study’s credibility deteriorated, limiting the therapeutic applicability of biomarkers. Like other biological markers, metabolomic biomarkers are hard to reproduce across studies due to demographic diversity and sample sources, variable experimental techniques, parameter settings in the metabolomics data, and biological changes in metabolite turnover rates (S. Huang et al. 2016). Overall, these constraints have hampered the introduction of novel cancer biomarkers into clinical practice. Due to system biology and bioinformatics tools, the integration of transcriptomics data with metabolomic profiling techniques has been employed in cancer research. It may provide more insight into these domains than either strategy alone (Ren et al., 2015). This novel technique might pave the door for new ways to explore pathogenesis via a system biology viewpoint and boost the reliability of biomarkers. However, no study has attempted to investigate CC in depth using metabolomics combined with transcriptomics in large samples.

Yang and his team conducted comprehensive metabolomics and transcriptomics-based investigation to examine specific dysregulated pathways and develop effective biomarkers for CC. They postulated that the genes and metabolites, the same biological processes, are frequently dysregulated together (S. Huang et al., 2016; Bjerrum et al., 2014). Therefore, metabolome and transcriptome data might be utilized to validate possible diagnostic biomarkers. The association between the identified metabolites and genes was then further explored using pathway and network studies, significantly boosting the reliability of the results.

A total of 5 groups were identified by classifying metabolites based on their metabolic profile. LysoPC (17:0), bilirubin, n-oleoyl threonine, tetracosahexaenoic acid, and 12-hydroxydodecanoic acid were chosen as potential biomarkers for CC. These biomarkers have AUC, specificity (SP), and sensitivity (SE), values of 0.99, 0.99, and 0.98, respectively (Yang et al. 2017).

MetaboAnalyst 3.0 was employed to perform pathway analysis on the sixty-two metabolites between CC patients and healthy controls. A total of thirty-one pathways were enriched, with 7 significant. These 7 pathways comprised glyoxylate and dicarboxylate metabolism, fatty acid biosynthesis, lysine biosynthesis, lysine degradation, histidine metabolism, steroid hormone biosynthesis, and citrate cycle. These pathways were primarily engaged in lipid metabolism (fatty acid and steroid hormone biosynthesis), amino acid metabolism (lysine biosynthesis and degradation, histidine metabolism), and carbohydrate metabolism (glyoxylate, dicarboxylate metabolism, citrate cycle,) all of which were significant in tumour tissue development and spread. For example, the up-regulated L-thyroxine was engaged in tyrosine and down-regulation of metabolites associated with the fatty acid metabolism and citrate cycle, leading to quick but inefficient energy metabolism. Proteins, membrane lipids, fatty acids, ATP, and nucleotides were required by the rapidly growing number of cells, explaining why metabolites involved in these processes were down-regulated.

A Global and targeted metabolomic profiling method (Khan et al., 2019) was utilized to discover plasma metabolite indicators and biomarkers for the non-intrusive prognosis of CC and risk factors associated with tumour development. CC causes differences in body metabolism and circulating metabolites. Investigating CC-related metabolic pathways might thus give valuable insights into the many processes used during tumour formation and progression and new approaches for early detection. While performing the global metabolite profile, they discovered that twenty-eight metabolites were substantially changed in CIN, cervical malignancies, and normal individuals. Thirty-seven normal and CC, twenty-nine CIN1 and CC, and thirty-three normal + CIN1 and CIN2/3 + CC pathways were effectively mapped. The metabolic pathways of aspartate, alanine, glutamate metabolism, proline and arginine metabolism, hypotaurine and taurine metabolism, and pyruvate metabolism were considerably changed between the control group and cancer patients, according to their pathway analysis results. In addition, 7 metabolites were identified to be significantly different in cervical carcinogenesis development after the validation stage. AUC (>0.80) and hierarchical cluster analysis (HCA) analysis were used to select them as possible biomarkers. Moreover, when paired with targeted altered metabolites in the validation stage, HPV, a recognized cause of CC advancement, was demonstrated to pose a significant risk.

In numerous investigations, metabolomics to identify CC biomarkers has shown to be quite promising. Khan and his colleagues reported that the analysis of circulating aspartate, hypoxanthine, glutamate, lactate, AMP, pyroglutamate, and proline levels might be used to differentiate healthy people from individuals with CIN and CC. In situations where metabolite levels were higher, the risk of getting CC increased. It was also shown that patients with elevated metabolite amounts and affirmative HPV infection had a greater chance of developing precancerous lesions and aggressive malignancy. However, more research is necessary to completely comprehend the link between metabolic alterations and CC development and their diagnostic potential.

## Artificial Intelligence/ Deep learning and CNN-based Approaches in CC

Machine learning and computational approaches are useful in drug discovery, and it has been well known since the 1990s (Chen et al., 2018). Radiotherapy, Brachytherapy, is a treatment that takes much time and extensive labour and shows variation. Novel studies are undergoing to delineate the benefits of speeding up the process and quality of the contours. Radiation and chemotherapy forms are the main guidelines for patients with IB2 and metastasis of CC, according to FIGO (International Federation of Gynecology and Obstetrics), several treatment strategies have been implemented, and one of them is neoadjuvant treatment has gained popularity in debulking of tumours, histology, tumour size, resectioning, etc. (Papadatos et al. 2015). Computed tomography, Magnetic resonance imaging, Positron emission tomography, and 18F -fluorodeoxyglucose are all very important in stage differentiating and therapeutic approaches (Herrera and Prior 2013). Radiomics is also one of the fastest-growing areas. It is vital in imaging analysis; the images’ high dimensional features of tumours, cysts, necrotic areas, and tumour heterogeneity can be visualized as marked through radiomics (Jajodia et al., 2021). This branch of computational biology with mathematical modelling from medical images gains prediction for survival and prognosis of cancer. In a recent study on CC that endeavours the knowledge through diffusion-weighted MRI and diffusion-derived quantitative parameters for imaging such as diffusion coefficient, radiomics are used to predict clinical outcomes. Moreover, lymph node metastasis and recurrence of CC have been predicted (Jajodia et al., 2021).

In a retrospective study of 462 cervical patients, extensive research was conducted by scientists, and researchers performed experiments to evaluate the performance of RA-CTVNet (Accurately delineating clinical target volumes) to speed up the contouring process and quality. Moreover, organ segmentation, auto segmentation based on the delineation of clinical target volume algorithms, and deep learning methods showed improvement in reducing the efforts of clinicians, therefore, increasing the accuracy and evaluation of the prognosis of CC (J. Shi et al. 2021).

## Therapeutic targets

Oncology is personalized medicine with advanced genetic tools and medications that produce better results. The therapeutic and treatment modalities are combined regimens for patients, such as platinum-based chemotherapy, radiation, radiotherapy, brachytherapy, etc. There are few biomarkers and inhibitors that work for certain therapies. For most CC patients with stage IVB and recurrent cases, palliative chemotherapy treatment is given, which is not manageable to curative treatment (Crowley, O’Cearbhaill, and Collins 2021). The overall median survival rate of persistent or recurrent cases and stage IV B is very low, with approximately 16-20% reported in the United States SEER N institute 2020 (Tian et al. 2020). There has been a fast development in cancer’s genetic and molecular profiles in recent years. (Table 5) (Figure 2)

**Table 5:**
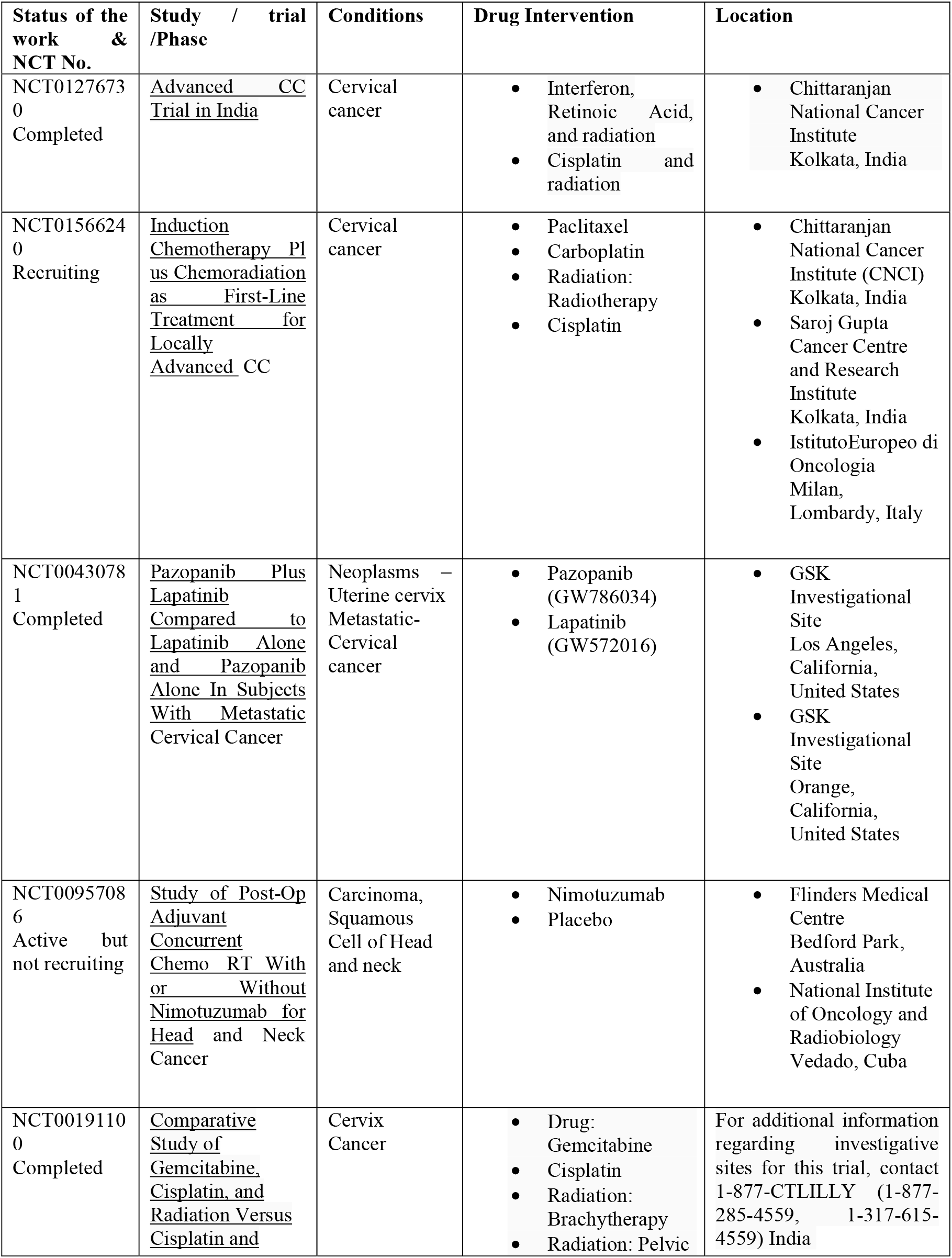
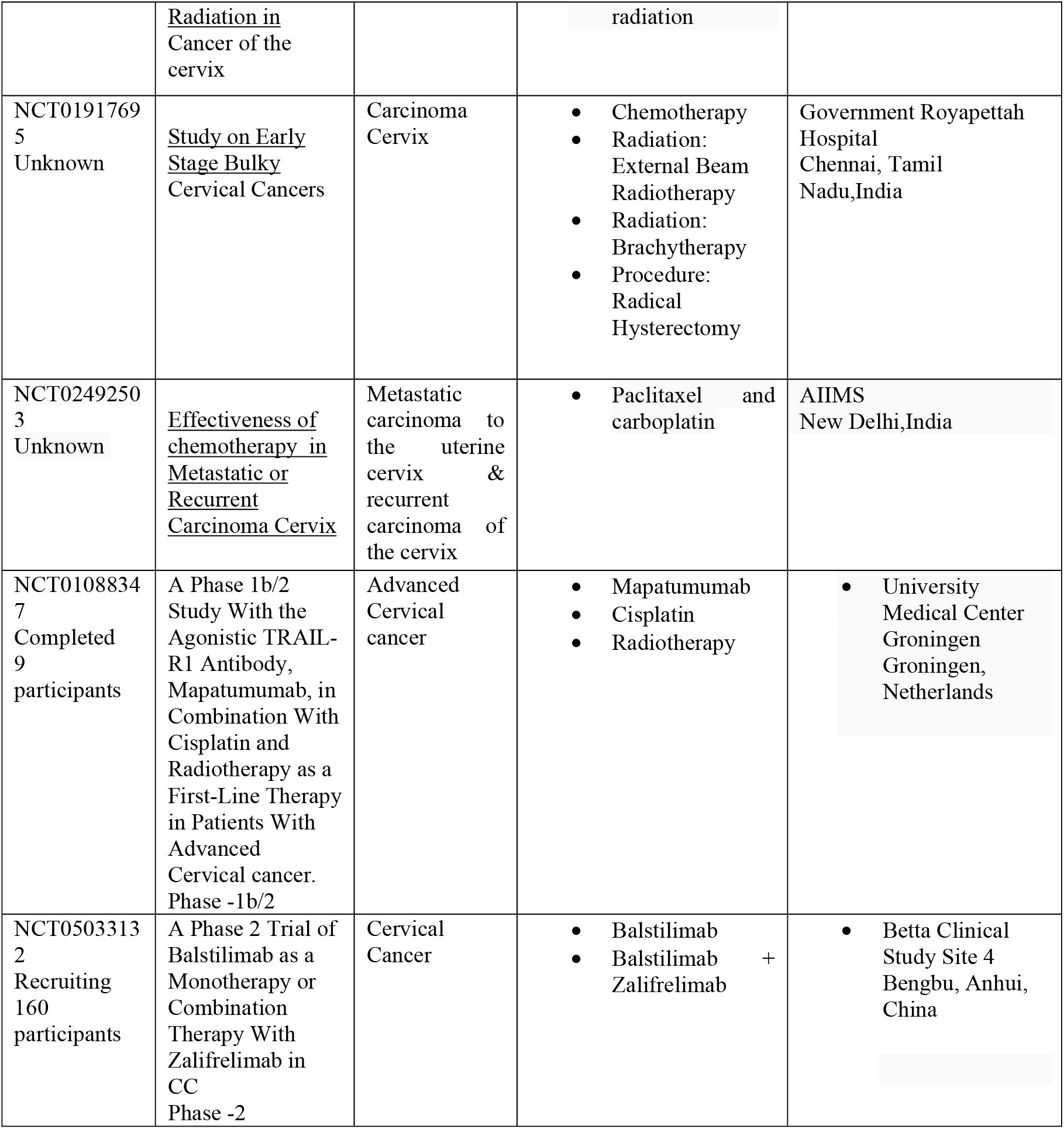
A table on Cervical Cancer Chemotherapeutic agents and trials. (www.gov.trials)

| Status of the work & NCT No. | Study / trial /Phase | Conditions | Drug Intervention | Location |
| --- | --- | --- | --- | --- |
| NCT01276730<br>Completed | <u>Advanced CC Trial in India</u> | Cervical cancer | Interferon, Retinoic Acid, and radiation<br>Cisplatin and radiation | Chittaranjan National Cancer Institute Kolkata, India |
| NCT01566240<br>Recruiting | <u>Induction Chemotherapy Plus Chemoradiation as First-Line Treatment for Locally Advanced CC</u> | Cervical cancer | Paclitaxel<br>Carboplatin<br>Radiation: Radiotherapy<br>Cisplatin | Chittaranjan National Cancer Institute (CNCI) Kolkata, India<br>Saroj Gupta Cancer Centre and Research Institute Kolkata, India<br>Istituto Europeo di Oncologia Milan, Lombardy, Italy |
| NCT00430781<br>Completed | <u>Pazopanib Plus Lapatinib Compared to Lapatinib Alone and Pazopanib Alone In Subjects With Metastatic Cervical Cancer</u> | Neoplasms – Uterine cervix Metastatic- Cervical cancer | Pazopanib (GW786034)<br>Lapatinib (GW572016) | GSK Investigational Site Los Angeles, California, United States<br>GSK Investigational Site Orange, California, United States |
| NCT00957086<br>Active but not recruiting | <u>Study of Post-Op Adjuvant Concurrent Chemo RT With or Without Nimotuzumab for Head and Neck Cancer</u> | Carcinoma, Squamous Cell of Head and neck | Nimotuzumab<br>Placebo | Flinders Medical Centre Bedford Park, Australia<br>National Institute of Oncology and Radiobiology Vedado, Cuba |
| NCT00191100<br>Completed | <u>Comparative Study of Gemcitabine, Cisplatin, and Radiation Versus Cisplatin and</u> | Cervix Cancer | Drug: Gemcitabine<br>Cisplatin<br>Radiation: Brachytherapy<br>Radiation: Pelvic | For additional information regarding investigative sites for this trial, contact 1-877-CTLILLY (1-877-285-4559, 1-317-615-4559) India |
|  | <u>Radiation in Cancer of the cervix</u> |  | radiation |  |
| NCT01917695<br>Unknown | <u>Study on Early Stage Bulky Cervical Cancers</u> | Carcinoma Cervix | • Chemotherapy<br>• Radiation: External Beam Radiotherapy<br>• Radiation: Brachytherapy<br>• Procedure: Radical Hysterectomy | Government Royapettah Hospital<br>Chennai, Tamil Nadu, India |
| NCT02492503<br>Unknown | <u>Effectiveness of chemotherapy in Metastatic or Recurrent Carcinoma Cervix</u> | Metastatic carcinoma to the uterine cervix & recurrent carcinoma of the cervix | • Paclitaxel and carboplatin | AIIMS New Delhi, India |
| NCT01088347<br>Completed<br>9 participants | A Phase 1b/2 Study With the Agonistic TRAIL-R1 Antibody, Mapatumumab, in Combination With Cisplatin and Radiotherapy as a First-Line Therapy in Patients With Advanced Cervical cancer. Phase -1b/2 | Advanced Cervical cancer | • Mapatumumab<br>• Cisplatin<br>• Radiotherapy | • University Medical Center Groningen Groningen, Netherlands |
| NCT05033132<br>Recruiting<br>160 participants | A Phase 2 Trial of Balstilimab as a Monotherapy or Combination Therapy With Zalifrelimab in CC Phase -2 | Cervical Cancer | • Balstilimab<br>• Balstilimab + Zalifrelimab | • Betta Clinical Study Site 4 Bengbu, Anhui, China |

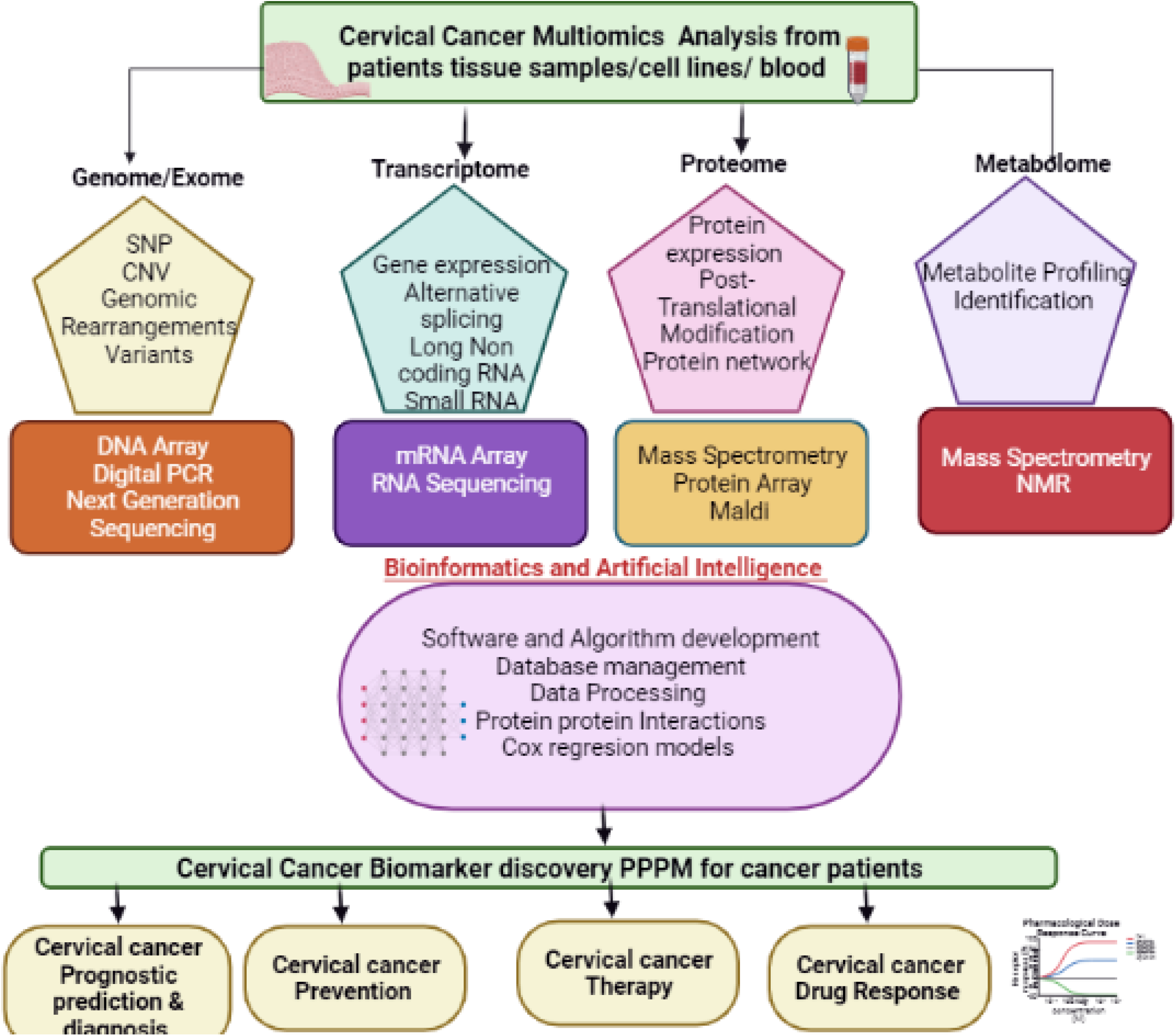

## Advantages of Multiomics

Multiomics data provides enormous advantages for investigating biological and molecular pathways more elaborately. It gives detailed information on how a genotype changes to a phenotype that undertakes the molecular transcripts and protein pathways to regulate the mechanism. Hallmarks of cancers allow the conversion of normal cells to invasive cancer. It includes specific phenotypic and genotypic molecular changes resulting in proliferation, angiogenesis, resisting cell death, and metastasis. Next-generation sequencing in cancer cells recognizes the mutations in pro-proliferative pathway genes and constitutes activation of signalling pathways but manifests in cells’ uncontrolled proliferation (Chakraborty et al. 2018). Multi-Omics in CC showed the clinical value of C1QC+ SPP1+, and TAM gene signatures subgrouped CC patients with various tumour stages and prognosis due to immune cell infiltration (X. Li et al. 2021). Through Genomics study Identification of SNPs, variants give information for early prognosis, diagnosis, prevention, and treatment in CC.

- High throughput provides accuracy at the transcriptomic level and digital gene expression level.

In proteomics study, while performing with 2D-DIGE provides quantitative comparative analysis using single gel, removing electrophoretic steps such as destaining and fixing (Karahalil 2016)

## Limitations

- NGS studies, PCR, and techniques for sequencing give results; however, DNA fragments of sequencing data should be verified with the Sanger sequencing technique in multi-omics research.
- It is difficult for transcriptome data to interpret and evaluate the statistics for large amounts of data (Karahalil 2016).
- Highly expensive
- In proteomics & metabolomics studies, there is a lack of proper information on scanning co-elution and sensitivity.

## Conclusion

Over the last decade, various large-scale multi-omics CC-based studies involving genomics, transcriptomics, proteomics, and metabolomics have demonstrated the highest contribution to protein abundance. Therefore, investigating CC-related metabolic pathways might give valuable insights into many processes used during tumour formation and progression and new approaches for early detection. However, according to researchers, multi-Omics technologies have enormous advantages and limitations to past studies. Therefore, it is better to opt for one or two omics technology at once to overcome the challenges. In addition, it is important to eliminate errors and contamination while performing NGS and better interpret and integrate the data meticulously.

## Abbreviations

CC: Cervical cancer
HPV: Human papillomavirus
CIN: Cervical Intraepithelial Neoplasia
FFPE: formalin-fixed paraffin mediated
SSE: stratified squamous epithelium
SCJ: Squamocolumnar Junction
SCCC: Squamous cell carcinoma of the cervix
NACT: Non-adjuvant chemotherapy
TOFMS: Time-of-flight mass spectrometry
BGLAP: Bone gamma-carboxy glutamic acid-containing protein
eEF1A1: Eukaryotic elongation factor 1-alpha-1
FIGO: International Federation of Gynaecology and Obstetrics
AUC: Area Under Curve
NGS: Next Generation Sequencing
NACT: Neoadjuvant Chemotherapy
CNN: Convolutional Neural Network
NMR: Nuclear magnetic resonance
MALDI - TOF: Matrix-Assisted Laser Desorption/Ionization-Time of Flight
MS: Mass Spectrometry

## Ethics approval and consent to participate

N/A

## Human and Animal rights

N/A

## Acknowledgement

We gratefully acknowledge the support from Lovely Professional University, Jalandhar, Phagwara, Punjab. Further, we thank the Department of Genetics, Osmania University Hyderabad, Telangana, for their generous support and the Bioclues Organisation for their support and contribution.

## Authors Contribution

**Sugunakar Vuree, Smita C Pawar**: Conceptualization, Conceived the idea, Proof Reading, **Santosh Kumari Duppala:**Wrote major part of the Manuscript, Abstract, Introduction, Clinicopathology of CC, Cervical precursor lesions histologic and cytologic classification, management of CC, Overview of Multiomics approaches for disease, Genomic studies of CC, Epigenomics, Artificial Intelligence/ Deep learning & CNN approaches, Therapeutic drugs, advantages of Multiomics, Limitations, Conclusion, **Rajesh Yadala**: Transcriptomics, **Ayushi Velingker:** Metabolomic, **Prashanth Suravajhala**: Critical Analysis, Proofreading of the manuscript.

